# DEGRONOPEDIA - a web server for proteome-wide inspection of degrons

**DOI:** 10.1101/2022.05.19.492622

**Authors:** Natalia A. Szulc, Filip Stefaniak, Małgorzata Piechota, Andrea Cappannini, Janusz M. Bujnicki, Wojciech Pokrzywa

## Abstract

The ubiquitin-proteasome system is a proteolytic pathway that removes damaged and unwanted proteins. Their selective turnover is initiated by ubiquitin (Ub) attachment, mainly by Ub ligases that recognize substrates through their short linear motifs termed degrons. A degradation-targeting degron comprises a nearby Ub-modified residue and an intrinsically disordered region (IDR) involved in interaction with the proteasome. Degron-signaling has been studied over the last decades, yet there are no resources for systematic screening of degron sites to facilitate studies on their biological significance, such as targeted protein degradation approaches. To bridge this gap, we developed DEGRONOPEDIA, a web server that allows exploration of degron motifs in the proteomes of seven model organisms and maps these data to Lys, Cys, Thr, and Ser residues that can undergo ubiquitination and to IDRs proximal to them, both in sequence and structure. The server also reports the post-translational modifications and pathogenic mutations within the degron and its flanking regions, as these can modulate the degron’s accessibility. Degrons often occur at the amino or carboxyl end of a protein substrate, acting as initiators of the N-/C-degron pathway, respectively. Therefore, since they may appear following the protease cleavage, DEGRONOPEDIA simulate sequence nicking based on experimental data and theoretical predictions and screen for emerging degron motifs. Moreover, we implemented machine learning to predict the stability of the N-/C-termini, facilitating the identification of substrates of the N-/C-degron pathways. We are confident that our tool will stimulate research on degron-signaling providing output information in a ready-to-validate context. DEGRONOPEDIA can be freely accessed at degronopedia.com.

## INTRODUCTION

Cellular differentiation and development, stress conditions, and environmental factors constantly challenge the integrity of the proteome in every eukaryotic cell. Maintaining protein homeostasis (proteostasis) requires the degradation of damaged or unwanted proteins and plays a crucial role in cellular function, organismal growth, and, ultimately, cell and organism viability (Douglas & Dillin, 2010; Morimoto & Cuervo, 2014). The ubiquitin-proteasome system (UPS) is a principal proteolytic component of the cellular proteostasis network (Glickman & Ciechanover, 2002). Enzymes operating within the UPS recognize protein substrates destined for degradation and label them by attaching a small, evolutionarily conserved protein ubiquitin, primarily to internal lysine residues (Kerscher et al., 2006). It is noteworthy that a growing number of evidence indicates that cysteine and serine/threonine residues can also function as ubiquitination sites forming thioester or hydroxyester bonds with ubiquitin, respectively (Kravtsova-Ivantsiv & Ciechanover, 2012; McClellan et al., 2019). Ubiquitination is mediated by an enzymatic cascade involving a ubiquitin-activating enzyme (E1), which activates the C-terminal glycine of ubiquitin and then transfers it to a ubiquitin-conjugating enzyme (E2). Subsequently, with ubiquitin ligase (E3) participation, Ub is transferred on the lysine residue of the substrate protein, forming an isopeptide bond with it (Glickman & Ciechanover, 2002). The proteasome complex recognizes ubiquitinated proteins and, through proteolysis, degrades them into short peptides that can be further processed (Komander & Rape, 2012).

E3 substrates can be targeted for degradation by exposing peptide signal motifs, called degrons, involving a lysine (or multiple lysines) acting as a ubiquitination site (Ravid & Hochstrasser, 2008; Varshavsky, 2019). Degrons comprise mainly short linear motifs, several amino acids long, and are thought to occur preferentially in disordered regions of proteins. Degrons can be constitutive, promoting continuous protein degradation, or conditional, emerging after post-translational modifications such as phosphorylation (Holt,2012)or after protease cleavage (Dissmeyer et al., 2018; Varshavsky, 2019). While degrons can be located anywhere in the protein sequence, those at the amino or carboxyl end, which are the initiators of the N- or C-degron pathway, respectively, have been the focus of extensive research over the last three decades (Bachmair et al., 1986; S.-J. Chen et al.,2017; Gonda et al., 1989; Hwang et al., 2010; Koren et al., 2018; Román-Hernández et al.,2009; Tasaki et al., 2005; Timms et al., 2019; Timms & Koren, 2020; Varshavsky, 2019; Yeh et al., 2021).

It is important to keep in mind that the recognition of a short linear motif by an E3 enzyme, followed by ubiquitination, may not be sufficient to lead to protein degradation. Guharoy *et al*. suggested that the short linear degron motif acts as a primary degron in the postulated tripartite degron architecture (Fig 1) (Guharoy et al., 2016). In this tripartite degron model, the secondary degron refers to lysine residues to which ubiquitin may be attached, and the tertiary degron indicates the flexible, intrinsically disordered region (IDR) in close proximity to the secondary degron, acting as a site to initiate protein unfolding prior to the entry into the proteasome. The secondary and tertiary degrons are suggested to play subsidiary roles that affect ubiquitin-signaling; the lack of a component of the tripartite degron model, e.g., an IDR near a ubiquitinated lysine, can result in non-proteolytic ubiquitination functions. Mutations leading to substitutions in degron motifs, as well as their secondary and tertiary degron sites, can therefore alter protein stability, contributing to diseases such as cancer and neurodegeneration (Eldeeb et al., 2022; Mészáros et al., 2017;Tokheim et al., 2021).

**Figure 1.**
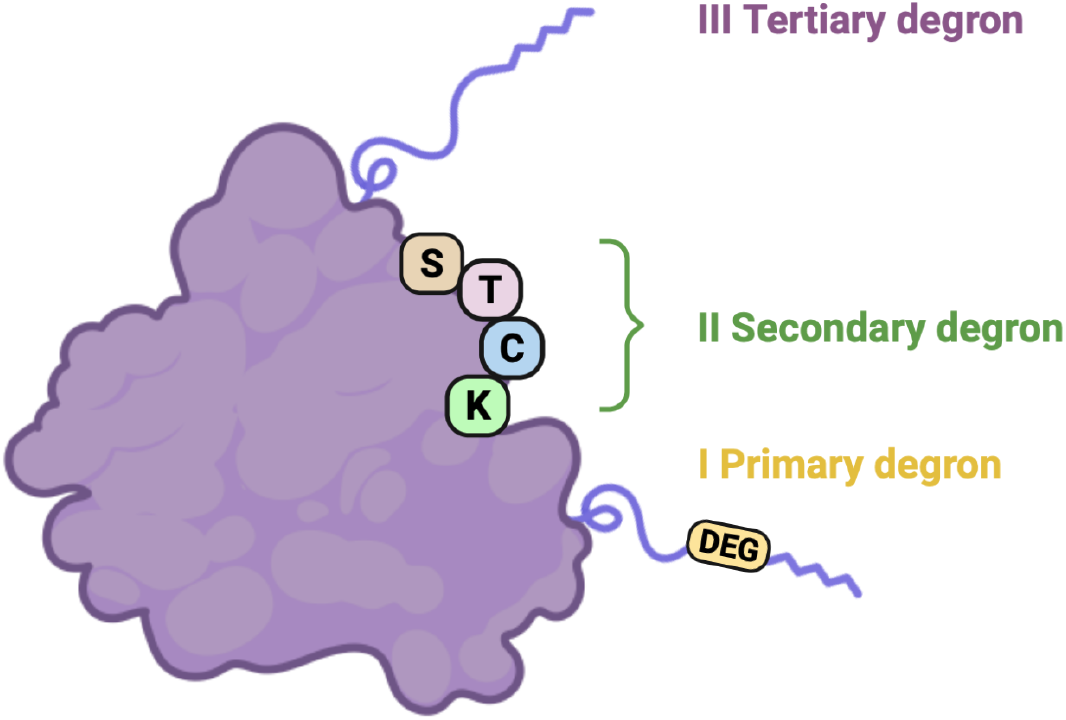
The tripartite degron model. The primary degron is a short linear motif recognized by the E3 ligase, localized preferentially within an IDR region of the protein. The secondary degron is a residue nearby the primary degron onto which ubiquitin transfer can occur (in our implementation, it is not only lysine (K) since ubiquitination can occur on cysteine (C), serine (S), or threonine (T)). The tertiary degron is an IDR close to the secondary degron, which acts as an unfolding seed initiating proteasome-dependent protein degradation. Modified from Guharoy et al., 2016, created in BioRender.com.

To better understand the specificity of the UPS and its deregulation in disease, degron sequences need to be discovered and matched with their respective degradation pathways. Recently, systematic approaches such as high-throughput experimental techniques, state-of-the-art proteomics technologies, and computational tools have been developed to understand the UPS selectivity (Coyaud et al., 2015; De Cesare et al., 2021;Kats et al., 2018; Koren et al., 2018; Nie et al., 2020; Timms et al., 2019; Wang et al., 2022;Yoshida et al., 2015). Biochemical and structural approaches complement the former to broaden our understanding of molecular mechanisms underpinning substrate selection by dedicated E3 ligases (X. Chen et al., 2021; Z. Chen et al., 2020; Chrustowicz et al., 2022;Yan et al., 2021). However, as the research on degron motifs and their physiological function is accelerating, there are few bioinformatics tools that would allow for degron motifs screening or analyzing. The anaphase-promoting complex/cyclosome (APC/C) degron repository (He et al., 2013) provides data on the sequence determinants of the three major classes of APC/C degron and overlays it with, e.g., disordered regions and post-translational modifications (PTMs). The eukaryotic linear motif (ELM) resource (Kumar et al., 2019) enables, among other functional sites, to detect 29 different degron motifs in the query protein and provides their structural context. A list of degron motifs, based on the data derived from the ELM resource and supplemented by manual curation, was released in 2017 as an interactive web table (dosztanyi.web.elte.hu/CANCER/DEGRON/TP.html, described in Mészáros et al., 2017). Finally, a deep learning model, deepDegron, was developed to predict degron disruption by mutations (Tokheim et al., 2021). However, deepDegron works as a standalone tool with specific input requirements - it accepts a list of mutations in a Mutation Annotation Format (MAF) file, containing particular columns with information on the gene of interest and its variants. To the best of our knowledge, no resource collects all known degron motifs and enables their systematic screening in query proteins, providing data on the degron site context, such as the postulated tripartite degron model.

Here we present DEGRONOPEDIA, the first web server designed to screen for known degron motifs in protein sequence or structure and provide their comprehensive sequence and spatial context complying with the tripartite degron mode. If the query protein comes from one of the selected model organisms - *A. thaliana*, *S. cerevisiae*, *C. elegans*, *D. melanogaster*, *D. rerio*, *M. musculus*, or *H. sapiens*, information on PTMs and known pathogenic mutations within the degron site and its neighborhood is also included. In addition, our tool allows the user to simulate the sequence cleavage by a number of different proteases to report N-/C-degron motifs in the newly emerged protein termini that may be of physiological interest. Our tool also allows users to examine their sequences and structures of interest for degron motifs. Moreover, DEGRONOPEDIA uses pre-trained machine learning (ML) models to predict the N-/C-terminal stability of the query protein. DEGRONOPEDIA is available free of charge in the form of a user-friendly web server at degronopedia.com.

## MATERIALS AND METHODS

### Inputs

Three input types can be used for querying DEGRONOPEDIA: (i) a UniProt ID of a protein from the reference proteome (according to the UniProt database (UniProt Consortium,2021)) of one of the selected model organisms - *A. thaliana*, *S. cerevisiae*, *C. elegans*, *D. melanogaster*, *D. rerio*, *M. musculus*, or *H. sapiens*, (ii) a protein sequence in the FASTA format, and (iii) a protein structure in the PDB format. The query protein must have between 50 and 8000 canonical amino acids regardless of input type.

#### Query by PDB

The submitted PDB file must not exceed 5MB size. Importantly, its B-factor column must carry either pLLDT (predicted Local Distance Difference Test; ranges 0-100; default for the AlphaFold models (Jumper et al., 2021)) or LLDT scores (Local Distance Difference Test; ranges 0-1; possible to obtain when predicting the model with RoseTTAFold (Baek et al., 2021), but requires further processing). The server does not support experimental PDB files since they lack long IDRs, which are crucial from the tripartite degron perspective. The detailed guide on the structure input type is available on the web server’s Tutorial web page.

### Customizable parameters

Screening for known degron motifs is dynamically performed on the web server, considering the eight user-customizable parameters. In short, they describe thresholds related to the distinct regions of the tripartite degron architecture and the structural models regarding the minimum values to reckon residues as buried or disordered (Table 1). A detailed description of the parameters and the appropriate visualizations are available on the web server’s Tutorial web page.

**Table 1.** Description of the eight user-customizable parameters available in the DEGRONOPEDIA.

| Parameter | Unit | Default value | Allowed values | Description |
| --- | --- | --- | --- | --- |
| **Primary degron-related** |  |  |  |  |
| 1. Degron flanking region in sequence | aa | 20 | 5-40 | Maximum sequence distance to regions upstream and downstream of a degron site to be considered flanking |
| 2. Degron flanking region in structure | Å | 20 | 5-40 | Maximum structural distance to residues near a degron site to be considered flanking (such residues are not necessarily close in sequence to the degron site) |
| 3. Region length to calculate degron disorder | aa | 10 | 1-20 | Maximum sequence distance to regions upstream and downstream of a degron site to be included in the degron mean disorder score based on pLLDT/LDDT values |
| **Secondary degron-related** |  |  |  |  |
| 4. Region length to calculate K/C/T/S disorder | aa | 3 | 1-15 | Maximum sequence distance to regions upstream and downstream of secondary degron (K/C/T/S) to be included in the secondary degron mean disorder score based on pLLDT/LDDT values |
| **Tertiary degron-related** |  |  |  |  |
| 5. Minimum IDR distance from K/C/T/S | aa | 10 | 5-40 | Minimum sequence distance of the secondary degron (K/C/T/S) to the continuous IDR region of defined length (see parameter 7) to consider it as a tertiary degron |
| **Structure-related** |  |  |  |  |
| 6. Minimum continuous IDR length | aa | 10 | 5-40 | Minimum number of subsequent (in sequence) disordered residues to be considered as IDR |
| 7. pLDDT/LDDT disorder threshold | % | 70 | 40-90 | Minimum value to recognize a residue as disordered based on its pLLDT/LDDT score |
| 8. Buried residue threshold | % | 20 | 5-60 | Minimum value to recognize the residue as buried based on its Relative Solvent Accessibility (RSA) |

### Implementation

We describe the calculations workflow for the query by UniProt ID since it provides the most exhaustive result information. The other query types are processed identically, with the exception that they do not cover certain analyses available when querying by UniProt ID as they do not access any additional data, i.e., structure model if submitting a sequence, PTMs or mutations.

#### General

The queried protein from the selected reference proteome is screened for the presence of each degron motif (see Degron motifs in the Datasets section), considering the defined degron position separately; N-termini and C-termini degrons are matched to the beginning or end of the sequence, respectively, whereas degron motifs classified as internal are searched in the entire sequence. The Gravy hydrophobicity index (Kyte & Doolittle,1982) of the first/last 15 amino acids of the queried protein is also calculated (Hickey et al.,2021; Kats et al., 2018). Upon data availability, the server provides the experimentally measured Protein Stability Index (PSI) of the N-/C-terminus and E3 ubiquitin ligases known to interact with the query protein (see N-/C-termini stability data and E3 interactome data in the Datasets section).

#### Structural data

Solvent-accessibility, location within an IDR region, and secondary structure are derived from the corresponding AlphaFold model. Of note, the AlphaFold models cover in its B-factor column pLLDT scores on a scale from 0 to 100, which estimate the accuracy of the modeled residues. Those with pLDDT above 70 are generally expected to be modeled well, while pLLDT below 70 correlates with disordered regions (Tunyasuvunakool et al., 2021). The server calculates (i) the IDR regions’ positions (based on the pLLDT threshold as defined in parameter 7 (Table 1) and the IDR minimum continuous length as defined in parameter 6 (Table 1)), (ii) the secondary structure, and (iii) Relative Solvent Accessibility (RSA) of each residue (based on the threshold as defined in parameter 8 (Table 1)). The secondary structure and solvent accessibility are calculated using the mkdssp software (Joosten et al., 2011; Kabsch & Sander, 1983), with the latter being normalized to the RSA based on the Sander method (Rost & Sander, 1994). The server further maps these data on each found degron motif and calculates the degron’s mean disorder (see parameter 3 in Table 1).

#### PTMs and mutations

As PTMs provide valuable information on potential degron modulation, the server maps the positions of known PTMs to each found degron motif and its flanking regions (regarding proximity in both sequence and structure, as defined in parameters 1 and 2 (Table 1), respectively). Moreover, the server also maps pathogenic missense mutations (only for the human proteins) to each degron motif and PTMs within its flanking regions.

#### Tripartite degron model context

The DEGRONOPEDIA server calculates the tripartite degron model for each of the found degron motifs. It searches for all potentially-ubiquitinated residues (in our implementation, these are not only lysines but also cysteines, serines, or threonines; K/C/T/S) within the degron flanking regions (regarding proximity in both sequence and structure, as defined in parameters 1 and 2 (Table 1), respectively) and maps their positions to the solvent accessibility and secondary structure data, location within an IDR, PTMs and pathogenic mutations; the server also calculates their mean disorder (see parameter 4 in Table 1). Finally, our tool reports the closest IDR (as defined in parameter 5 in Table 1) and its distance (both in sequence and structure) to each of the aforementioned secondary degrons.

#### Protein cleavage

The DEGRONOPEDIA server simulates the cleavage of queried protein based on the experimentally-validated proteolytic sites derived from the MEROPS database as well as predictions of cleavage sites for 35 different proteolytic enzymes using the Pyteomics Python module (Goloborodko et al., 2013; Levitsky et al., 2019), which implements the cleavage prediction rules of the PeptideCutter Expasy web server (Gasteiger et al., 2005). Next, our tool screens each newly emerged N-/C-terminus for known degron motifs as described before.

### N-/C-terminus stability predictions

For a protein sequence or structure submission, the user may optionally request predictions of its N-/C-terminus stability using our pre-trained ML models.

#### ML models development

The experimental N-/C-termini stability data, expressed as Protein Stability Index (PSI) values measured for 23-mers covering N-/C-termini of the nearly complete human proteome (Koren et al., 2018; Timms et al., 2019), were used to develop the ML models. Of note, from the N-termini dataset, we considered only peptides’ variants without the first methionine residue. Although methionine is the first amino acid incorporated into each new protein, it is not always the first amino acid in mature proteins undergoing post-translational removal (Varland et al., 2015; Yeom et al., 2017). Therefore, we decided to train an ML model to predict N-terminal stability without the initial methionine. The datasets were split into the training and testing set (in the ratio of 90:10), and each testing set remained untouched until the final testing of the models. Predictive models were built using the CatBoost regressor (Prokhorenkova et al., 2018) and trained on the aforementioned datasets, separately for each terminus. Hyperparameters of the CatBoost were optimized with the Optuna framework (Akiba et al., 2019) with five-fold cross-validation (using random permutations cross-validation implemented in scikit-learn Python library, with 20% validation set; (Pedregosa et al., 2012)). Descriptors used for building the models include the sequence of the peptide, RDKit descriptors (RDKit: Open-source cheminformatics; http://www.rdkit.org), Gravy hydrophobicity index (Kyte & Doolittle, 1982), and Peptides module (its Python version; github.com/althonos/peptides.py; (Osorio et al., 2015)). All descriptors were calculated for the whole sequence and the first (excluding the N-terminal methionine) or the last (for the C-terminus) ten, eight, six, four, and two amino acids. The performance of the final models was evaluated using the testing set and an *R^2^* coefficient, reaching the values of 0.815 for the C-terminus and 0.796 for the N-terminus (Fig 2). To visualize the predicted PSI N-/C-terminus value, the server maps it to the distribution of the corresponding experimental N-/C-termini stability dataset and classifies it as unstable/moderately unstable/average/moderately stable/stable, denoted by quantile thresholds of 0.2/0.4/0.6/0.8/1.0.

**Figure 2.**
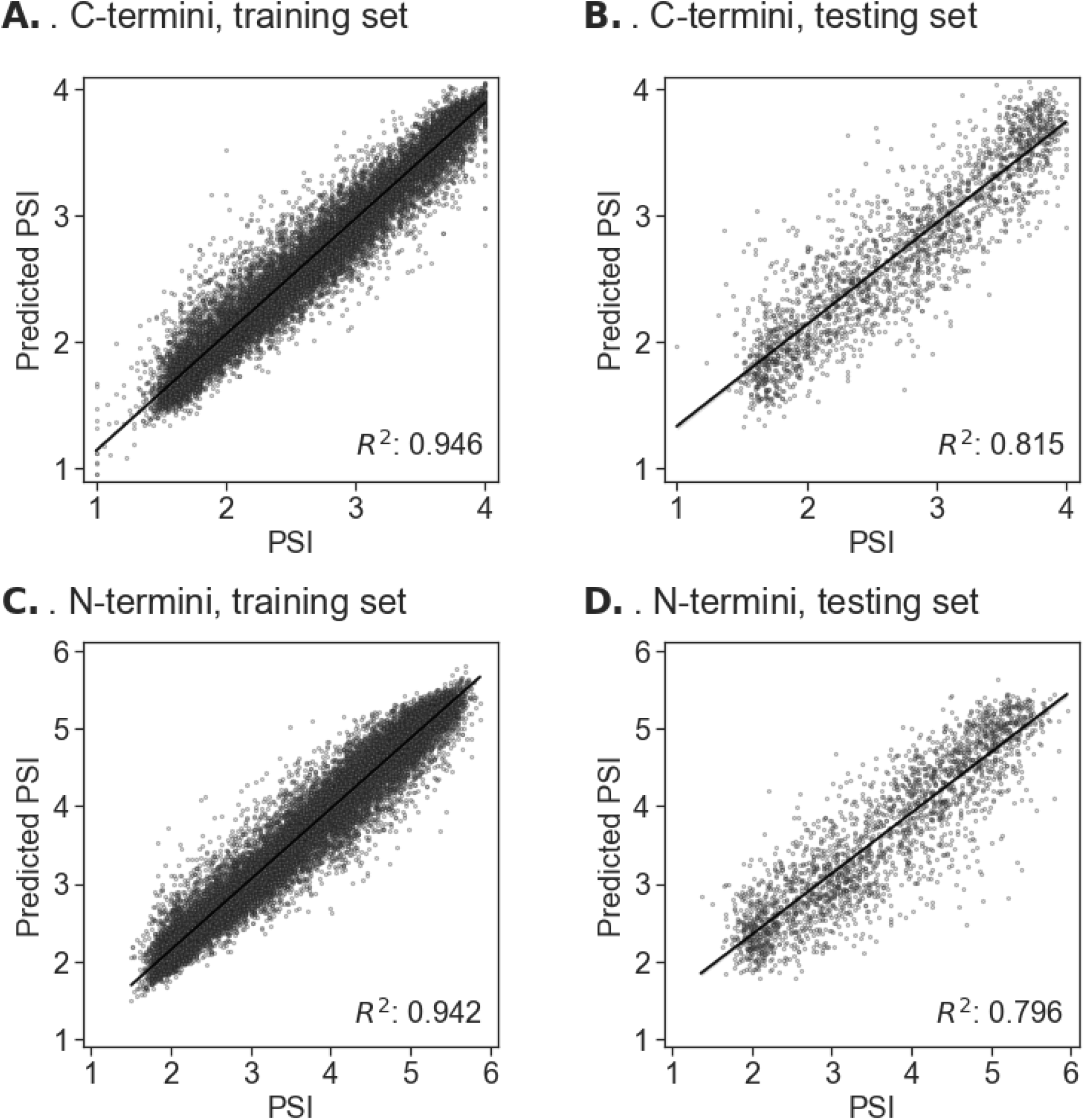
Predictions of the PSI using ML CatBoost regression models for C-termini (A, B) and N-termini (C, D); for the training (A, C) and testing set (B, D). Scatter plots show a regression line with a 95% confidence.

### Outputs

The output information may be downloaded as a xlsx file, with corresponding data saved to separate sheets.

### Visualization

The Feature-Viewer tool (Paladin et al., 2020) was used for visualizations.

### Datasets

All the described datasets, except for the Degron motifs dataset, are applicable only when querying by UniProt ID.

#### Degron motifs

Over 400 degron motifs were obtained from the literature (X. Chen et al.,2021; Guharoy et al., 2016; Koren et al., 2018; Maurer et al., 2016; Timms et al., 2019;Varshavsky, 2019; Yan et al., 2021). Each motif was defined as either N-terminus, C-terminus, or internal, regarding its occurrence location. Moreover, additional data, including organisms in which the degron motif was found, degron type, known E3 ligases recognizing it, and subsidiary information, were added upon availability. N-terminus (Timms et al., 2019) and C-terminus (Koren et al., 2018) degron motifs derived from Global Protein Stability assays were selected based on their delta PSI (Protein Stability Index) value defined as >0.7 and >0.3, respectively.

#### Reference proteome data

Reference proteomes for *A. thaliana*, *S. cerevisiae*, *C. elegans*, *D. melanogaster*, *D. rerio*, *M. musculus,* and *H. sapiens* were obtained from the UniProt database (IDs: UP000006548, UP000002311, UP000001940, UP000000803, UP000000437, UP000000589, UP000005640, respectively). Sequences shorter than 50 and longer than 8000 amino acids were excluded from further analysis as very short peptides may not contain all the components of the tripartite degron model, and extremely long sequences are sporadic and require computationally exhaustive calculations. Where literature data were available, relevant information about each protein, i.e., its validated degron motifs or ubiquitinated lysines leading to proteasome-dependent degradation, was added.

#### Structural data

For each proteome as described above, its appropriate structural models were downloaded from the AlphaFold Protein Structure database (Varadi et al., 2022), excluding models for proteins exceeding the above-mentioned sequence length thresholds. As AlphaFold models of proteins longer than 2700 amino acids are split to separate files (containing overlapping fragments of the model), an in-house script utilizing the Biopython module (Cock et al., 2009) was used in each such case to superimpose overlapping residues, merge the structures’ parts and save them to a single pdb file.

#### N-/C-termini stability data

The N-/C-termini stability data were obtained from the Global Protein Stability assays (Koren et al., 2018; Timms et al., 2019). These data covered the stability of N-/C-terminal 23-mers of 48251 (variants with N-terminal methionine and without it) and 22564 human proteins, respectively, represented as the Protein Stability Index (PSI), where its highest values (6 for the N-termini and 4 for the C-termini) indicate the most stable, thus possibly deprived of degron motifs, peptides. Consecutive mapping of N-/C-terminus PSI is performed by the server only for human proteins and based on the N-/C-terminal 23-mers identity, not the protein IDs. Similarily as in our ML models implementation (see N-/C-terminus stability predictions in Materials and methods), the N-termini peptides’ variants without the first methionine residue were considered, and thus, the reported N-terminal PSI value refers to the stability of 23-mer N-terminus with absent methionine.

#### Post-translational modification data

The post-translational modification datasets were obtained from the iPTMNet (Huang et al., 2018) (phosphorylation, acetylation, ubiquitination, methylation, N-Glycosylation, O-Glycosylation, C-Glycosylation, S-Glycosylation, sumoylation, myristoylation, S-Nitrosylation), PhosphoSitePlus (Hornbeck et al., 2015) (phosphorylation, acetylation, ubiquitination, methylation, O-Glycosylation, sumoylation) and the PLMD (Z. Liu et al., 2011, 2014; Xu et al., 2017) (neddylation and formylation) databases as well as from the literature, where we manually compiled datasets of non-canonical ubiquitination (Carvalho et al., 2007; Chua et al., 2019; Hensel et al., 2011; Ishikura et al.,2010; X. Liu & Subramani, 2013; Williams et al., 2007) and arginylation (Wong et al., 2007).

#### E3 interactome data

The complete interactomes were acquired from the BioGRID (Oughtred et al., 2021), IntAct (Orchard et al., 2014) and UbiNet 2.0 (Li et al., 2021) databases and from the literature, where we manually compiled a dataset of E3-substrate interactions absent in the aforementioned databases from (Koren et al., 2018; Oh et al.,2017; Weaver et al., 2017). As we provide information about the known interactions of the query protein with various E3 ligases (including components of their complexes), it was necessary to filter proteins from the selected model organisms annotated as E3 ligases. These annotations were obtained from the AMIGO web server (Ashburner et al., 2000;Carbon et al., 2009; Gene Ontology Consortium, 2021) by submitting the GO:0061630 query for each model organism and mapping the unique hits to their UniProt IDs. In the case of human E3 ligases, an additional annotation dataset was manually created, tabulating information from the AMIGO web server with ESBL (Medvar et al., 2016) and the UbiNet 2.0 data. Finally, the BioGRID and IntAct interactomes were accordingly filtered to derive only the E3-substrate interactions’ sub-datasets (the UbiNet 2.0 already contained the E3-substrate interactions only).

#### Mutation data

A dataset of human mutations classified as pathogenic was obtained from the COSMIC database (Tate et al., 2019). Among the available mutation types, only ‘Substitution - Missense’ mutations were considered, as they have the least disruptive effect on the entire protein compared, e.g., to the frameshift or nonstop mutations, and may most severely impact the degron motif itself.

#### Proteolytic cleavage sites data

The experimental proteolytic cleavage sites with adherent information about the involved proteolytic enzymes were derived from the MEROPS database (Rawlings et al., 2018). The MEROPS dataset was subsequently filtered to contain only cleavage sites classified as physiologically relevant with present information about their exact position in the sequence.

## RESULTS

The DEGRONOPEDIA web server accepts three input types: (i) UniProt ID of a protein from the reference proteome of one of the selected model organisms - *A. thaliana, S. cerevisiae, C. elegans*, *D. melanogaster*, *D. rerio*, *M. musculus*, or *H. sapiens*, (ii) protein sequence in the FASTA format, and (iii) protein structure in the PDB format. Depending on the input type, different granularity of degron-related information is provided (Fig 3), with the most comprehensive data available for the UniProt ID query. However, regardless of the submitted query type, the protein sequence is always screened for the presence of over 400 degron motifs, which were obtained from the literature (X. Chen et al., 2021; Guharoy et al.,2016; Koren et al., 2018; Maurer et al., 2016; Timms et al., 2019; Varshavsky, 2019; Yan et al., 2021).

**Figure 3.**
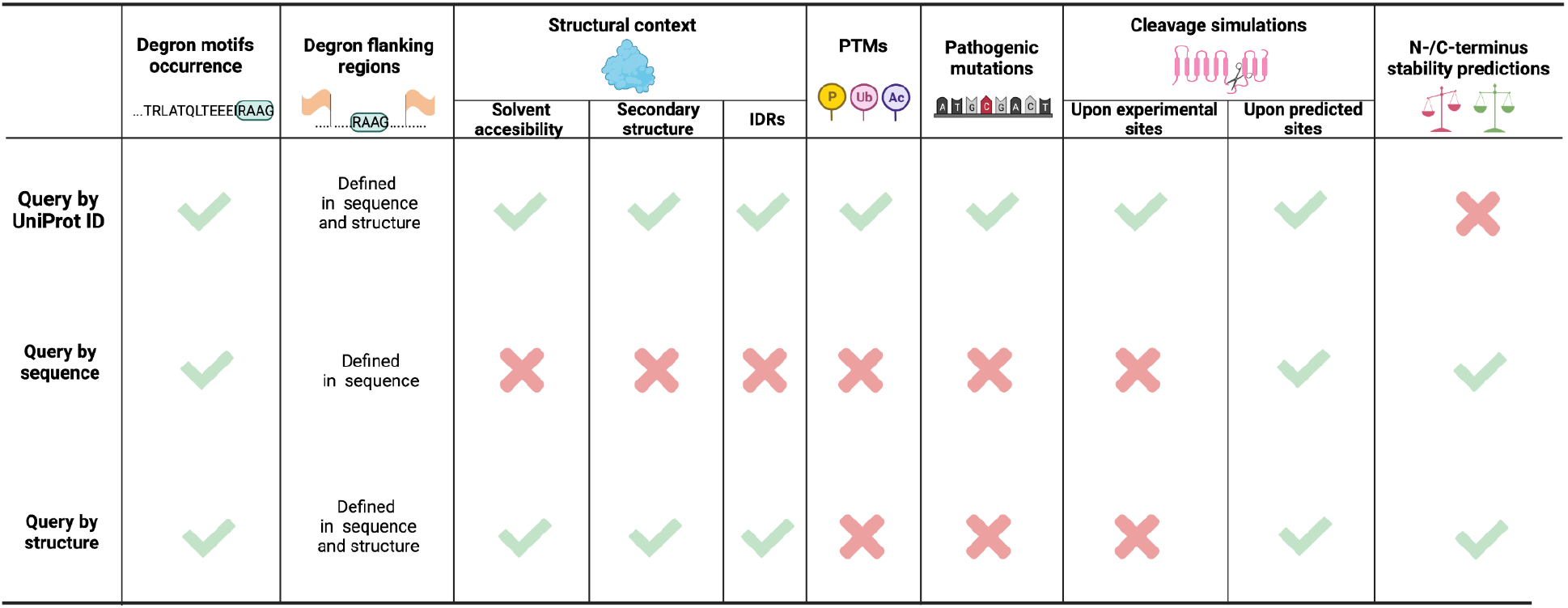
Comparison of the result information obtained upon different query types in the DEGRONOPEDIA server. Created in BioRender.com.

### Query by UniProt ID

The UniProt ID input type provides the most exhaustive information about the putative and experimentally-validated degron sites overlaid with additional information, i.e., solvent-accessibility, occurring PTMs or pathological mutations nearby. Of note, it is the only input type available in the DEGRONOPEDIA that provides experimentally-derived information on PTMs, mutations, physiologically-relevant proteolytic cleavage sites, and E3 interactors.

The queried protein is screened for the presence of degron motifs from the dataset prepared as described before. The Gravy hydrophobicity index (Kyte & Doolittle, 1982) of the N-/C-termini is also calculated, as hydrophobicity was shown to play an essential role in the N-/C-degron recognition by different E3 ubiquitin ligases (Hickey et al., 2021; Kats et al.,2018). The server also provides the information on N-/C-terminus stability (defined as Protein Stability Index (PSI); currently only for human proteins), collected from Global Protein Stability studies (Koren et al., 2018; Timms et al., 2019), and reports on E3 ubiquitin ligases interacting with the queried protein to provide further insights into possible degron-receptor interactions.

Since solvent-accessibility, location within an IDR region, or lack of secondary structure are important premises of a site acting as an actual degron, our tool derives such information from the corresponding AlphaFold model and overlays it on degron sites. The server also introduces degron flanking regions in sequence and structure to further extend the degron local context and maps the aforementioned features to these. PTMs are involved in multiple cellular processes, i.a., molecular interactions, protein folding, solubility, or signaling (Ramazi & Zahiri, 2021). Degron sites undergo various PTMs, with the primary role of phosphorylation (such degrons are often referred to as phospodegrons), which can modulate their exposure (Holt, 2012). Hence, the server reports PTMs (up to 14 types) occurring within each found degron motif and its flanking regions. Amino acid substitutions in degron motifs can lead to altered protein stability, contributing to severe diseases such as cancer (Mészáros et al., 2017) or neurodegeneration (Eldeeb et al., 2022), indicating critical sites for proper protein function. Therefore, the server provides information about known pathogenic missense mutations within the degron motifs and PTMs in their flanking regions.

For each of the found degron motifs (primary degron), its context to the secondary and tertiary degron is calculated according to the tripartite degron model postulated by Guharoy and colleagues (Fig 1) (Guharoy et al., 2016). In particular, the server provides information on solvent-accessibility, secondary structure, location within an IDR, mean disorder, PTMs, and pathogenic mutations for each potentially-ubiquitinated residue (secondary degron) located within the degron flanking regions. Finally, the server reports the closest IDR to each of the aforementioned secondary degrons.

It has been shown that protein turnover may be regulated by different proteolytic enzymes that cleave the protein, leading to new N- and C-termini which may act as degrons (Dissmeyer et al., 2018; Varshavsky, 2019). Therefore, the server simulates protein cleavage, after which it analyzes the newly emerged N- and C-termini for degron motifs.

All the results are reported in the form of comprehensive tables, where each column is described in detail on the web server’s Tutorial web page. Localization of degron motifs, coils, buried residues, IDRs, PTMs, and pathogenic mutations are mapped to the query sequence and visualized using the Feature-Viewer tool (Paladin et al., 2020). All the output information may be downloaded as a xlsx file, with corresponding data appropriately sorted to separate sheets.

### Query by sequence

Submitting a sequence provides the most limited output data compared to two other input types available in the DEGRONOPEDIA since it carries the least information about the protein. Briefly, the degron motifs screening and cleavage simulations (upon predicted proteolytic sites) are performed identically as described for the UniProt ID query. For the tripartite degron model representation, only the secondary degrons (located within the degron flanking region in sequence) are reported since no structural data is provided.

However, an additional feature of this query type is the possibility to run ML models to predict the stability, expressed as the PSI, of the N-/C-terminus of the submitted protein sequence. The predicted PSI is visualized as a publication-ready figure, since the server maps each predicted PSI to the distribution of its corresponding experimental N-/C-termini stability dataset and classifies it as unstable/moderately unstable/average/moderately stable/stable. We recommend running the N-/C-termini stability predictions only on proteins from higher mammals, as our ML models were trained on stability datasets of human proteins (see N-/C-terminus stability predictions in Materials and methods).

As sequence input yields restricted calculations and, consequently, limited output information, we encourage users to query by structure and submit a model of their protein of interest from state-of-the-art structure prediction tools such as AlphaFold or RoseTTAFold.

### Query by structure

The structure input type was designed as a compromise between the granularity of the output information and the possibility of analyzing proteins other than from the reference proteomes of selected model organisms. All the calculations are performed identically as when querying by the UniProt ID, except no data on PTMs, mutations, physiologically-relevant proteolytic cleavage sites, or E3 interactors are provided. The user may run our ML models to predict the stability of the N-/C-terminus, identically as when querying by sequence.

### Example application

As a case study, we chose the human p53 tumor suppressor protein (UniProt ID P04637), for which numerous experimental data on turnover are available. Proteasomal degradation of p53 is based on degron sequences and post-translational modifications (Asher et al.,2005; Melvin et al., 2016; Yang et al., 2006). Within the p53 protein, DEGRONOPEDIA annotated six degrons (we ran the calculations with the default parameters), one as of the N-terminus pathway and the rest as internal degrons. These include the FSDLWKLL motif (positions 19-26; also defined more broadly as F[^P]{3}W[^P]{2,3}[VIL]), which is recognized by the E3 ubiquitin ligase MDM2 (Böttger et al., 1997; Kussie et al., 1996; Schon et al.,2002). In addition, the server reported the presence of two sites (positions 248-256 and 338-346) corresponding to motifs recognized by the anaphase-promoting complex (APC/C) E3 ubiquitin ligase, which, to our knowledge, has not been previously described as engaged in p53 degradation. Interestingly, the server annotated that the putative degron recognized by APC/C (positions 338-346) is frequently mutated, which may indicate its functionality. Its neighboring lysines (positions 291-292, 351 (<20Å) and positions 319-321, 357 (<20 aa)) are subject to several PTMs, including ubiquitination, and thus, could represent a secondary part of the degron. Fig 4 shows how DEGRONOPEDIA summarizes found degron sites, plots PSI values of N-/C-termini on the experimental data distribution, and assigns secondary/tertiary degrons in the p53 sequence and structure.

**Figure 4.**
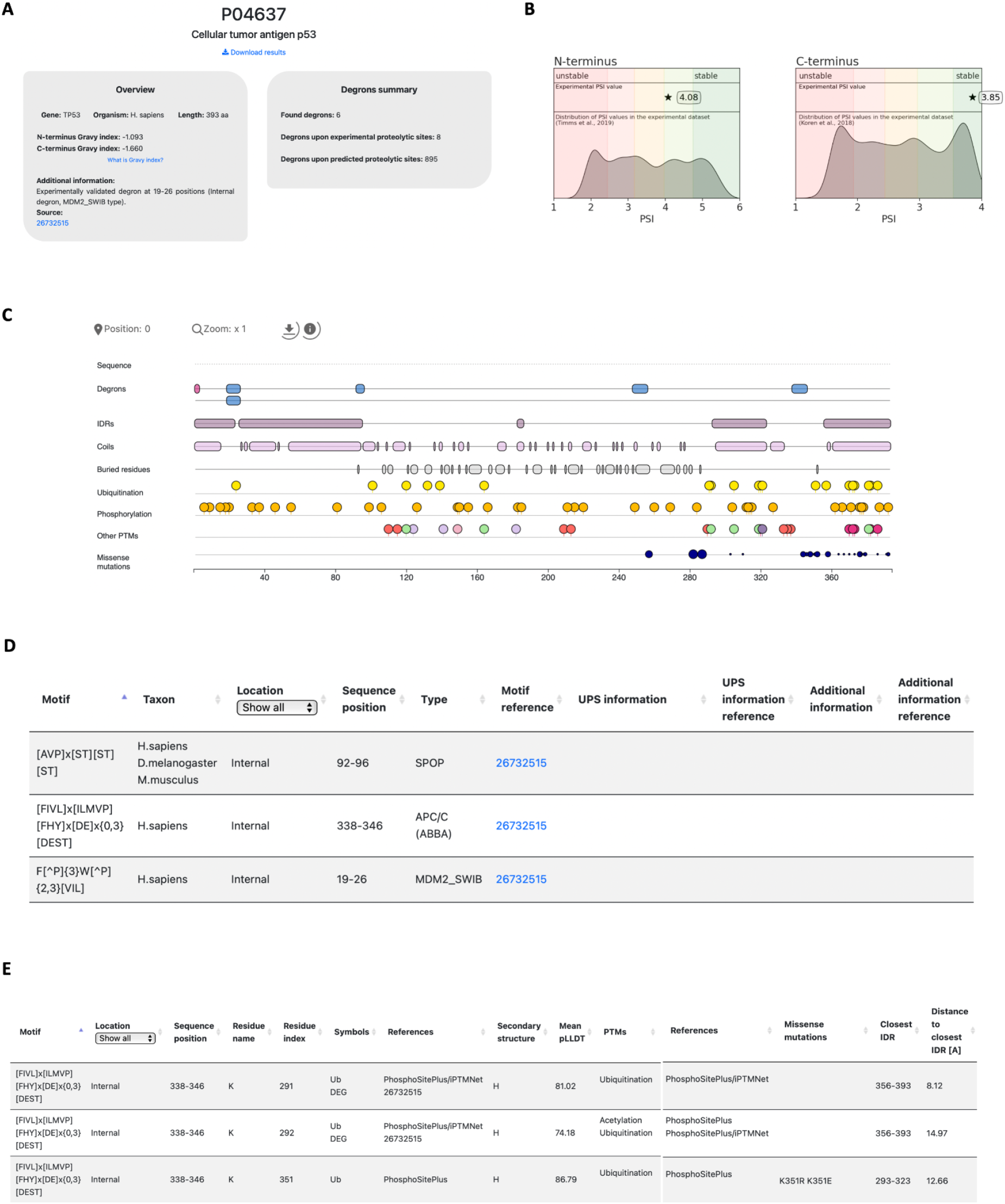
Overview of the DEGRONOPEDIA server on the example of p53 tumor suppressor protein. **(A)** overview panel with general information on the p53 protein (left) and degrons’ summary panel with information on the number of found degron motifs (right); **(B)** experimental PSI values of N-/C-terminus of p53 plotted on the distribution of experimental stability datasets; **(C)** visualization of the degron motifs, structural data, PTMs and mutations on the protein sequence; **(D)** table summarizing found degron motifs; **(E)** extract of table denoting the secondary and tertiary degrons associated with a putative degron motif at positions 338-346.

## DISCUSSION

DEGRONOPEDIA is a web server that integrates multifaceted degron screening in a query protein, complying with the postulated tripartite degron model by Guharoy and colleagues (Guharoy et al., 2016), with an intuitive interface and convenient visualization of the analysis results. It accepts three input types: UniProt ID, the sequence in the FASTA format, and structure in the PDB format, providing different levels of the output information depending on the submitted query type, with the most comprehensive results available for the UniProt ID query. The operability of our web server allows intuitive checking for the presence of known degron motifs in combination with overlapping PTMs and pathogenic mutations (if available) and detection of new N-/C-pathway degron motifs after simulated proteolysis. Additionally, DEGRONOPEDIA can predict protein N-/C-termini stability based on the submitted sequence or structure, utilizing the pre-trained ML models.

However, DEGRONOPEDIA has some limitations. Regardless of the input type, the protein sequence must be between 50 and 8000 amino acids long (which roughly corresponds to the maximum 5MB file size when querying by structure) and should contain the 20 canonical residues only. Of note, the query by UniProt ID option is currently available only for proteins belonging to the reference proteomes of the seven most studied model organisms. Thus, users who would like to analyze a protein from another taxon, or an isoform not included in the reference proteome, have to choose between the query by sequence or query by structure options. Another limitation is that the submitted structure must be a monomer and should contain valid pLLDT or LDDT scores, as this information is used to extract the position of IDRs. Therefore, since experimentally obtained structures do not hold these values, they cannot serve as inputs to DEGRONOPEDIA. We reasoned that their lack of long IDRs does not make them the input of choice from the tripartite degron model perspective, especially in the light of the currently available state-of-the-art tools such as AlphaFold or RoseTTAFold which provide high-quality complete protein models. However, one must keep in mind that deriving information on the disorder residues based on the pLLDT-defined threshold may be inaccurate. Aderinwale and colleagues recently analyzed the correlation between disorder predictions obtained from different disorder prediction software tools and the regions from AlphaFold models with pLDDT scores below 0.5 and 0.7 (Aderinwale et al., 2022). Their results showed that only 30-50% of residues with low pLLDT scores correspond to disordered regions predicted using other methods, while the rest would adopt folded structures. Therefore, one must not treat the pLLDT scores as an absolute metric for defining IDRs. For this reason, we provided a customizable parameter so the user may manipulate the pLLDT threshold below which the residue is considered disordered to partially circumvent this issue.

In future versions of the DEGRONOPEDIA, we plan to further enhance the accuracy of the disorder region predictions by allowing the user to upload their own list of disordered residues as well as we will provide the option to predict IDRs using a state-of-the-art software tool to derive the disorder consensus in concert with the pLLDT/LDDT scores. In addition, we aim to extend the functionality and customizability of the web server by providing an option to define own degron motifs to screen for in the query protein. In addition, we will implement a new analysis type allowing for consecutive N-/C-terminus depletion, which would serve as a guide in the site-directed mutagenesis studies aiming to delete the degron site and, simultaneously, not create a novel one. As visualizations provide the most understandable and user-friendly way to report the result data, we also plan to map the elements of the degron tripartite model on a structure viewer, allowing for its comprehensive inspection on the native protein surface. Last but not least, the detailed output provided by the DEGRONOPEDIA heavily depends on the literature data. Thus, we plan to integrate new datasets relevant to the degron-signaling upon newly published research on a rolling basis.

## DATA AVAILABILITY

The web server is available at degronopedia.com. This website is free and open to all users, and there is no login required.

## ACKNOWLEDGMENTS

We would like to acknowledge Martina Bevilacqua for her invaluable support regarding the requested enhancements of the Feature-Viewer tool. We would like to express our gratitude to Dr. Natalia Gumińska for preparing the visual identification of the DEGRONOPEDIA. We would like to thank Dr. Neil Rawlings for providing exhaustive explanations on the MEROPS database usage. We would also like to acknowledge current members of the Pokrzywa group, in particular, Lilla Biriczová and Dr. Abhishek Dubey.

This research was carried out in part with the support of the Interdisciplinary Centre for Mathematical and Computational Modelling (ICM) at the University of Warsaw under computational allocation no G88-1177 to F.S.

